# Beyond Sequence Similarity: ML-Powered Identification of pHLA Off-Targets for TCR-Mimic Antibodies Using High Throughput Binding Kinetics

**DOI:** 10.1101/2025.09.19.677256

**Authors:** Alexander Sinclair, Stefan Krämer, Christoph Reinhart, Jennifer Stehle, Simon Schuster, Tobias Herz, Hoor al Hasani, Pranav Hamde, Oliver Selinger, Joerg Birkenfeld

**Affiliations:** BioCopy GmbH, Emmendingen, Germany; BioCopy AG, Basel, Switzerland

**Keywords:** HLA, MHC, pHLA, pMHC, TCR-like antibody, peptide-MHC complex, X-Scan, MAGE-A4, off-target toxicity, preclinical de-risking, machine learning, binding kinetics, SCORE, BLI, T2-binding

## Abstract

T-cell receptor mimic (TCRm) antibodies are an emerging class of tumor-targeting agents used in advanced immunotherapies such as bispecific T-cell engagers and CAR-T cells. Unlike conventional antibodies, TCRms are designed to recognize peptide–human leukocyte antigen (pHLA) complexes that present intracellular tumor-derived peptides on the cell surface. Due to the typically low surface abundance and high sequence similarity of pHLAs, TCRms require high affinity and exceptional specificity to avoid off-target toxicity. Conventional methods for off-target identification such as sequence similarity searches, motif-based screening, and structural modelling focus on the peptide and are limited in detecting cross-reactive peptides with little or no sequence homology to the target. To address this gap, we developed EpiPredict, a TCRm-specific machine learning framework trained on high-throughput kinetic off-target screening data. EpiPredict learns an antibody-specific mapping from peptide sequence to binding strength, enabling prediction of interactions with unmeasured pHLA sequences, including sequence-dissimilar peptides. We applied EpiPredict to two distinct TCRms targeting the cancer-testis antigen MAGE-A4. The model successfully predicted multiple off-targets with minimal sequence similarity to the intended epitope, many of which were experimentally validated via T2 cell binding assays. These findings establish EpiPredict as a valuable tool for lead optimization of TCRms, enabling the identification of antibody-specific off-targets beyond the scope of traditional peptide-centric methods and supporting the preclinical de-risking of TCRm-based therapies.

## INTRODUCTION

Expanding the target space for antibody-based therapeutics remains a major goal in oncology, as the number of clinically validated surface targets remains limited. Most approved monoclonal antibodies recognize cell surface proteins, which comprise only 26% of the human proteome [1], inherently restricting the therapeutic landscape. The 2022 approval of Tebentafusp, a bispecific T cell receptor (TCR)-antibody fusion protein that redirects T cells to uveal melanoma via a peptide derived from the intracellular protein gp100 presented by HLA-A*02:01, provided clinical proof-of-concept that intracellular antigens can be exploited therapeutically through peptide Human Leukocyte Antigen (pHLA) complexes [2, 3]. This breakthrough has fuelled intense interest in pHLA-targeted therapies [4]. T-cell receptor mimic (TCRm) antibodies represent a promising modality within this space. By emulating the specificity of native TCRs, TCRms can bind intracellular-derived peptide antigens presented by HLA on the cell surface, significantly expanding the number of targetable proteins. Moreover, their compatibility with versatile antibody formats such as antibody-drug conjugates (ADCs), bispecific T-cell engagers (TCEs), and CAR-T constructs offers clear advantages in design flexibility, manufacturability, and therapeutic delivery [5, 6].

Despite their promise, developing therapeutic TCRms remains highly challenging. Target pHLA complexes are often present at very low surface densities, necessitating high-affinity binding, often in the picomolar range. However, because the TCRm binding interface typically focuses on a short peptide epitope of just 9-10 amino acids, achieving high affinity without compromising specificity is inherently difficult [7]. The limited sequence length and structural similarity among many pHLA complexes significantly raise the risk of off-target cross-reactivity. Such unintended interactions can result in severe toxicities, underscoring the need for rigorous off-target assessment during early development [8].

Conventional strategies for off-target prediction including sequence motif analysis, positional X-scans, peptide-HLA binding prediction, and structural modelling are largely peptide-centric [9]. While these tools identify pHLA variants with similar sequences, they often fail to detect off-target peptides with little or no sequence homology to the intended target that may still engage the TCRm paratope through shared structural or physicochemical features.

To address these limitations, we developed EpiPredict, a TCRm-centric machine learning (ML) framework for off-target identification. EpiPredict leverages high-throughput kinetic binding data from X-scan and pHLA microarrays to train antibody-specific neural networks that learn sequence-affinity relationships. This enables prediction of TCRm binding strength across the full proteome, including peptides with little or no sequence similarity to the intended target. Unlike conventional peptide-centric methods based on sequence similarity or structural heuristics, EpiPredict learns directly from empirical binding outcomes, allowing it to identify functionally relevant off-targets regardless of sequence conservation.

We applied EpiPredict to two distinct TCRm antibodies targeting the cancer-testis antigen MAGE-A4 [10]. The model accurately predicted diverse, antibody-specific off-target pHLA interactions, many of which were validated experimentally using label-free kinetic profiling and T2 cell-based flow cytometry assays. These results demonstrate the utility of EpiPredict as a predictive, structure-aware tool for de-risking TCRm candidates by uncovering off-target liabilities beyond the reach of traditional peptide-centric methods.

## RESULTS

### Overview of the EpiPredict Workflow

Our EpiPredict workflow consists of 5 main phases (Figure 1). In order to demonstrate the approach, we selected two TCRm antibodies (Antibody A and B) that should specifically target and bind the MAGE-A4 decapeptide (230GVYD-GREHTV239) within the HLA-A*02:01 context. During the course of the study, we extensively characterise the two TCRms and use the EpiPredict workflow to predict and validate additional off-targets, that might be physiologically critical.

**Figure 1.**
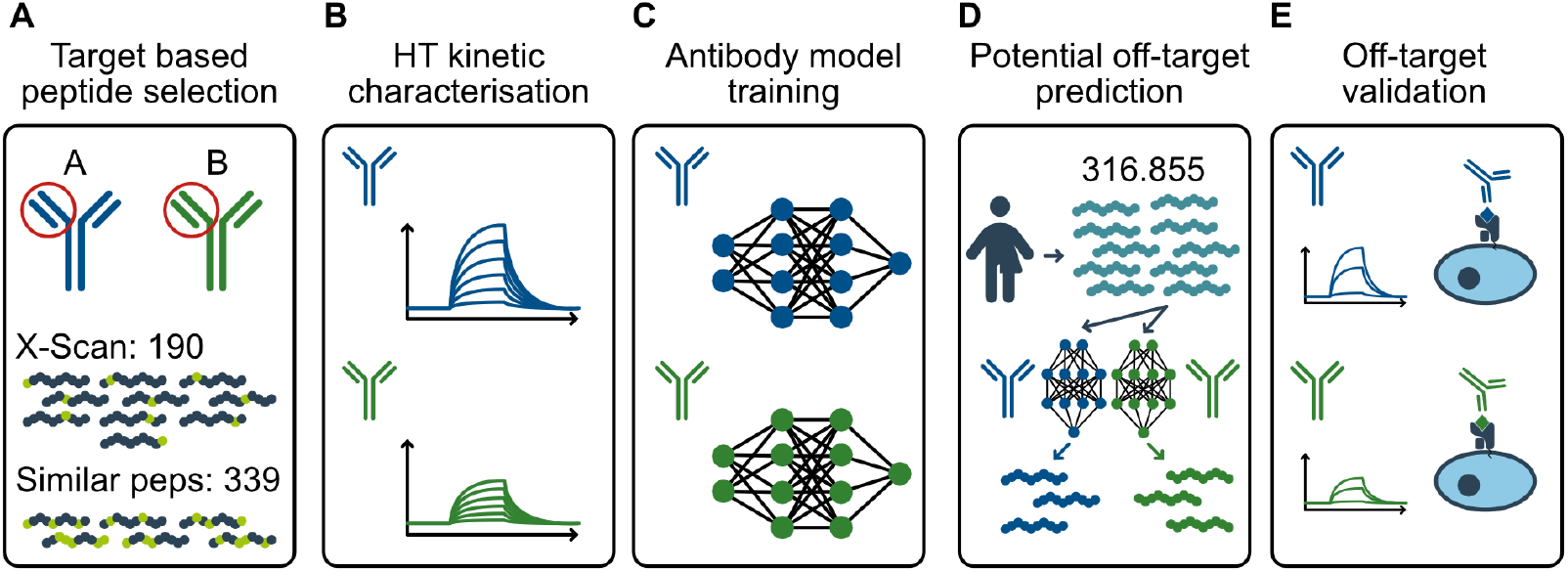
EpiPredict core workflow. (A) A dataset of 530 peptides (339 EpiTox-predicted off-targets and 190 X-scan variants) were selected. (B) Peptides were formatted as pHLA-A*02:01 microarrays and profiled using SCORE for each antibody. (C) Binding data was used to training machine learning models for each antibody. (D) 316,855 peptides from the human proteome likely to be presented by HLA-A*02:01 (NetMHCpan rank ≤2) were fed to the models. (E) High-probability hits prioritized for confirmatory in in-vitro assays.

A dual strategy was employed to create the initial biologically meaningful peptide set for the first round of antibody characterization (Figure 1 A). First, we generated a positional X-scan of the MAGE-A4 peptide, comprising all 190 single amino acid substitutions, to systematically explore TCRm sequence tolerance. Second, we applied a custom peptide-centric workflow, EpiTox (manuscript in preparation), which scans the human proteome with a position-specific similarity scoring matrix to identify sequences related to MAGE-A4. For this study, peptides with up to five substitutions were included. Candidates were further filtered for predicted HLA-A*02:01 binding (NetMHCpan [11]) and tissue-specific expression (Human Protein Atlas; [12]). This yielded 339 potential off-target peptides, which together with the 190 X-scan variants formed a dataset of 530 peptides. Thereafter, high-throughput, label-free interaction profiling was performed using SCORE (Single-Color Reflectometric Interference Detection). For each TCRm, kinetic parameters, association rate (*k*_*on*_), dissociation rate (*k*_*off*_), and equilibrium dissociation constant (*K*_*D*_), were measured across all 530 complexes. Complexes with measurable *K*_*D*_ values were classified as binders and all others were labeled as non-binders (Figure 1 B). Subsequently, the labeled data was used to train one model per antibody (Figure 1 C). The models were then applied to systematically predict the TCRm binding to an artificial library consisting of 316.855 theoretical human proteome-derived decapeptides (Figure 1D). All predicted high-probability peptides were experimentally validated using SCORE microarrays and T2 cell-based flow cytometry assays, completing the discovery-validation loop (Figure 1 E).

### Target based antibody characterization

The two sequence-distinct TCRm antibodies, A and B were first characterised using BLI and exhibited low-nanomolar affinities, Kd=6.1nM and Kd=4.5nM to their HLA-A*02:01–MAGE-A4 target (Figure 2 A & B). These findings were further supported by an X-scan peptide-HLA positional library kinetic measurement. Here, a peptide panel was generated by systematically substituting each residue of the WT MAGE-A4 decapeptide with all 19 alternative amino acids (X-SCAN). The two TCRms exhibited partially overlapping but distinct binding preferences for the MAGE-A4 peptide-HLA positional variants (Figure 2 C,D). Both antibodies show to require an arginine at position 6 for efficient binding. However, single substitutions at other positions had differential effects. For example, substitutions at position 9 with positively charged residues enhanced binding of antibody A but had no effect on antibody B. Together, the kinetic and positional scanning data suggest that both antibodies possess distinct paratope characteristics, which may mediate differential recognition of additional, unseen peptide–HLA complexes.

**Figure 2.**
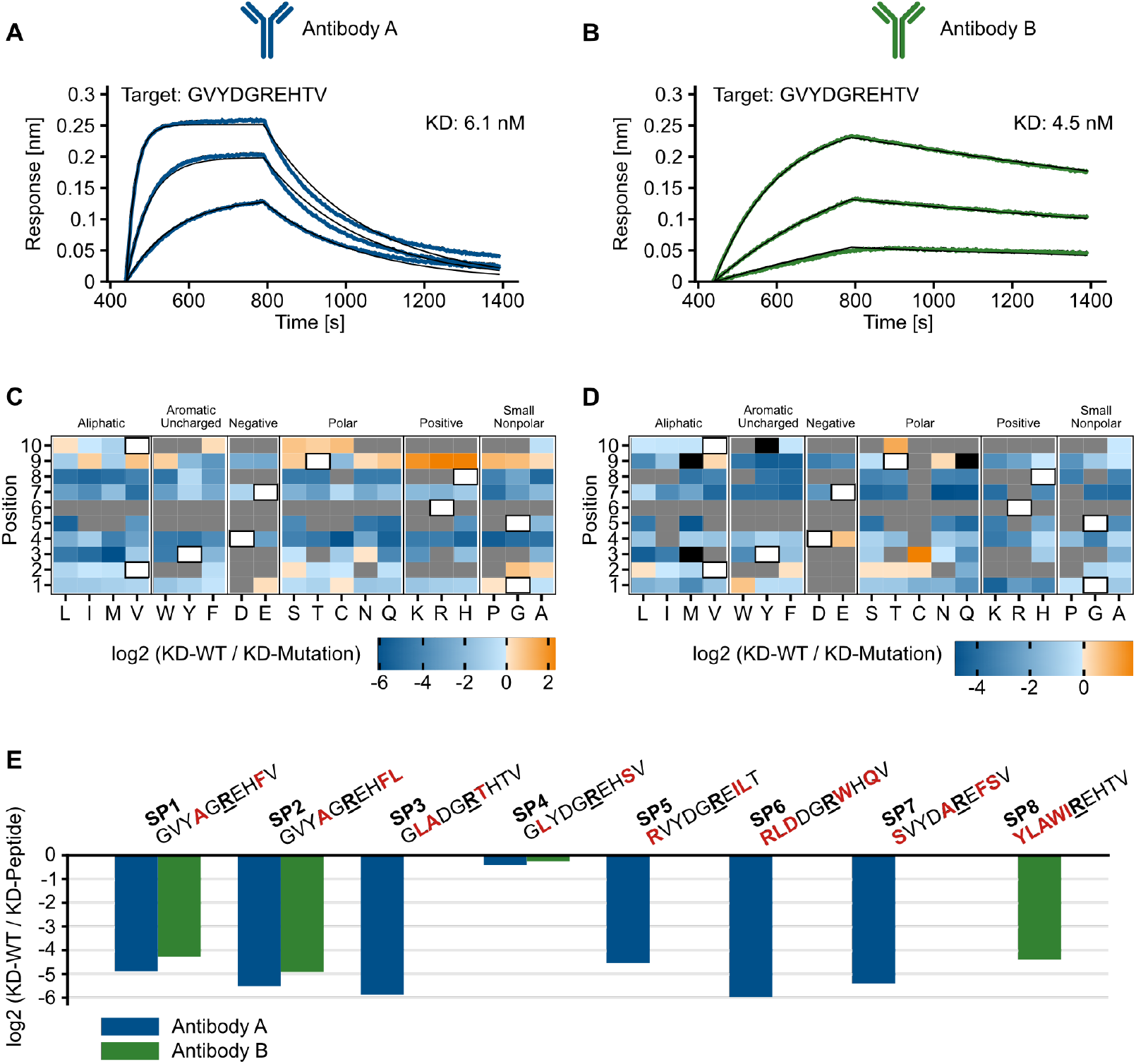
Kinetic profiling of two TCRms, Antibody A (A, C, E) and Antibody B (B, D, E). (A, B) Biolayer interferometry (BLI) binding kinetics of TCRms with target MAGE-A4. Association and dissociation phases are shown, with 1:1 global fit curve overlaid (solid black lines). (C, D) Heatmap of *K*_*D*_ ratio values from positional peptide library scanning (X-Scan) of the MAGE-A4 wild-type peptide (GVYDGREHTV). Higher log_2_(*K*_*D*_ WT/*K*_*D*_ mutant) ratios, indicative of stronger binding, are shown in warmer colors; lower ratios are shown in cooler colors. White tiles denote wild-type residues, grey tiles indicate positions where *K*_*D*_ values could not be determined consistent with abolished binding, and black tiles represent positions where peptide loading could not be confirmed by *β*_2_M staining. (E) log_2_(*K*_*D*_ WT/*K*_*D*_ mutant) ratios of peptides with more than one mutation that showed binding to either entibody A (blue) or antibody B (green). In this case, no bar is equal to no binding observed.

In addition to the X-Scan analysis, we also performed a high-throughput kinetic characterisaion of the two TCRms against 339 potential off-target peptides that had more than one mutation compared to the target motive (Figure 2 E). Antibody A showed binding against 7 and antibody B against 4 out of the 339 off-targets. The overlap was 3. In all but one cases, the binding was weaker (higher *K*_*D*_ value) compared to the target peptide. Interestingly, both antibodies bound SP4, a peptide derived from the MAGE-A8 protein, with affinities comparable to the WT peptide. In general, MAGE-A8 and MAGE-A4 share a high sequence identity and a their similar presentation pattern by HLA-A*02:01 is to be expected. Similarly, both TCRms recognized other MAGE-family peptides, including MAGE-C2 (SP1) and MAGE-A11 (SP”), albeit with a weaker affinity. The partial overlap of off-target binding suggest that the two TCRms engage the MAGE-A4-pHLA complex via distinct paratope conformations and interaction modes, supported by differences in affinity, CDR sequences, and structural frameworks.

### Antibody modeling and TCRm off-target predictions

The labeled kinetic interaction data from the previous section was used to follow a supervised machine learning approach. We trained independent neural networks for each TCRm using their respective binding datasets to predict the probability of TCRm-pHLA interactions for yet unseen peptides in the HLA-A*02:01 context. Our choosen model architecture is illustrated in Figure 3, A. The performance of the trained neural networks was evaluated using out-of-fold predictions from cross-validation training. Classification metrics were computed at a 50% probability threshold, with results summarized Table 1.

**Table 1.**
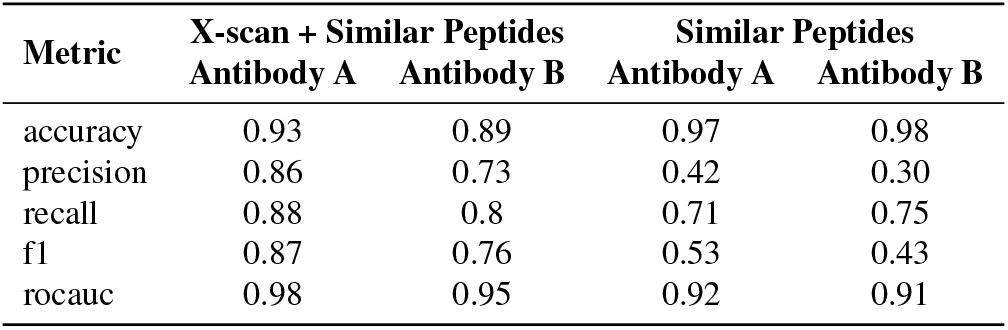
Performance Metrics Comparison.

**Figure 3.**
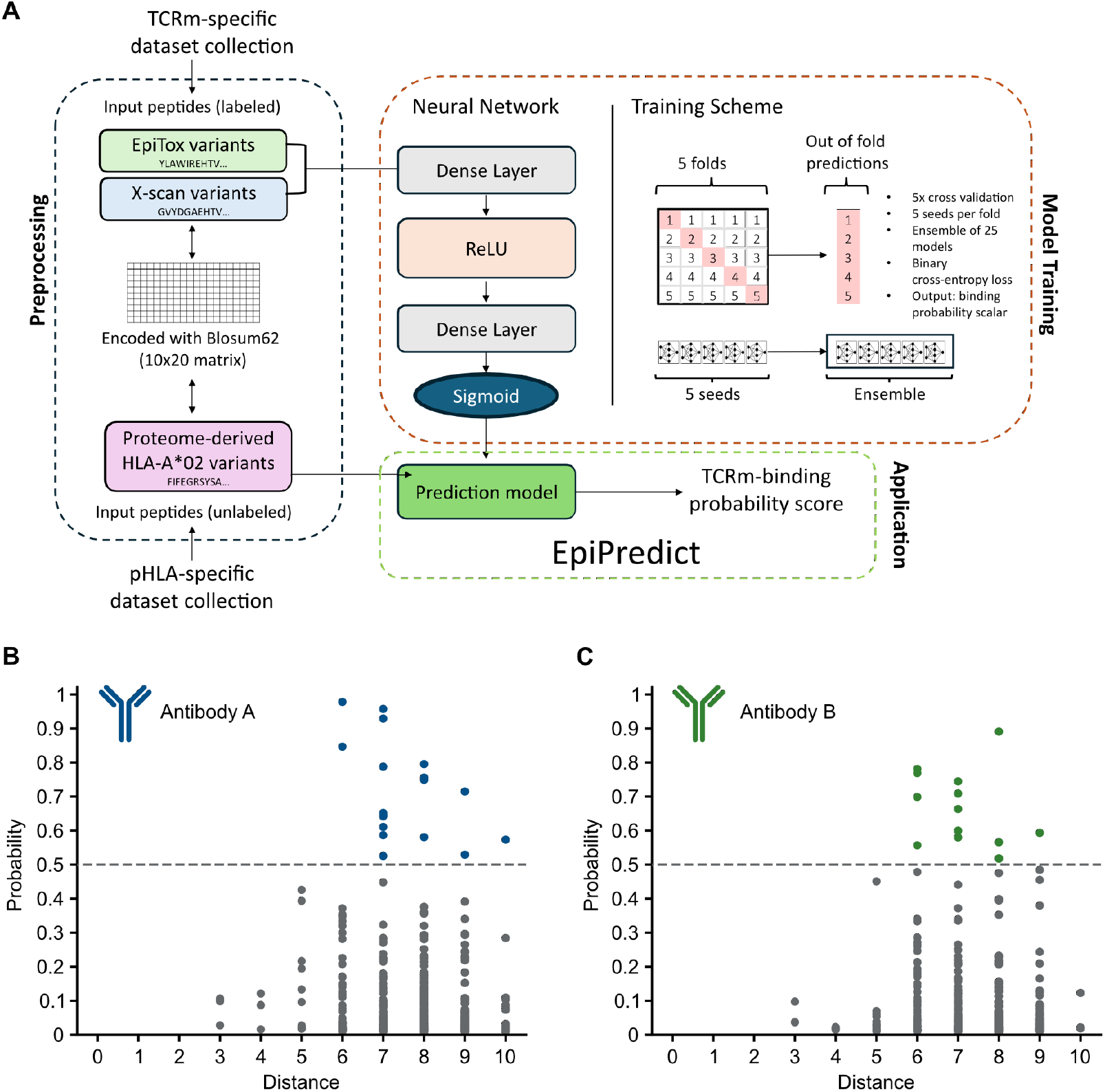
EpiPredict Machine learning model training and predictions. (A) Model architecture and training scheme. (B) & (C) Predicted antibody binding probabilities across sequence dissimilarity levels. Scatter plot of individual peptide-antibody pairs showing EpiPredict-predicted binding probabilities (y-axis) as a function of sequence dissimilarity to the MAGE-A4 decapeptide, quantified by Hamming distance (x-axis; 1–10 substitutions). Each point represents a unique peptide differing by the indicated number of amino acids. Peptides with predicted binding probability *>*0.5 were selected for experimental validation. The distribution demonstrates that predicted binders occur even among peptides with high sequence dissimilarity.

The models demonstrated high predictive performance, achieving accuracies ≥89 for both antibodies (Table 1). Over the entire dataset, including both X-Scan and similar peptides, the models achieved precision scores of 0.86 and 0.73 for Antibody A and Antibody B respectively, indicating strong ability to accurately distinguish between binders and non-binders.

However, when the analysis was restricted to only the similar peptides subset, a substantial decline in precision was observed despite maintained high accuracy. Precision dropped to 0.42 and 0.30 for antibody A and B respectively. This performance degradation can be attributed to the severe class imbalance present in the similar peptides, where only 7 out of 339 peptides (2.1%) were binders for Antibody A and 4 out of 339 peptides (1.2%) were binders for Antibody B. Despite this extreme imbalance, the models still demonstrated meaningful predictive power, predicting the majority of off-targets correctly with recall scores of 0.71 for antibody A and 0.75 for antibody B.

To validate model performance and identify highly dissimilar pHLAs with binding potential, we applied the models to a predefined decapeptide sequence space comprising 316,855 peptides derived from the full human proteome and restricted to HLA-A*02:01. A threshold probability of *>*50% was applied to extract potential off-target candidates from the models predictions. In total, 29 peptides were predicted as binders across both TCRms, with the majority being antibody-specific. This represents a remarkably selective binding profile, with only 0.009% of the total peptide space meeting the binding threshold, highlighting the stringent specificity of the TCRm models. Of these predicted binders, 18 peptides were predicted for antibody A (Figure 3 B) and 14 for antibody B (Figure 3 C). Three peptides were predicted to bind both. Also, we observed that several peptides with high predicted binding scores exhibited substantial divergence from the WT sequence.

### Experimental Validation of Potential Off-Target Peptides

To experimentally validate the predicted off-targets, we performed SCORE based kinetic interaction analysis. Here, we pooled the predicted peptides for both antibodies and used the combined set for the validation for both antibodies. For antibody A, 5 of the 29 predicted peptides exhibited measurable binding, with dissociation constants in the high to mid-nanomolar range (Figure 4, A). In contrast, antibody B only showed detectable binding against two peptides (PP5 and PP9) of the 29 predicted candidates (Figure 4, B). Notably, these peptides were also validated for antibody A, suggesting overlapping specificity. Sequence analysis of the validated binders highlighted the critical importance of arginine at position 6. All other peptides lacking this residue failed to bind which is consistent with the prior X-scan results (Figure 2, C, D).

**Figure 4.**
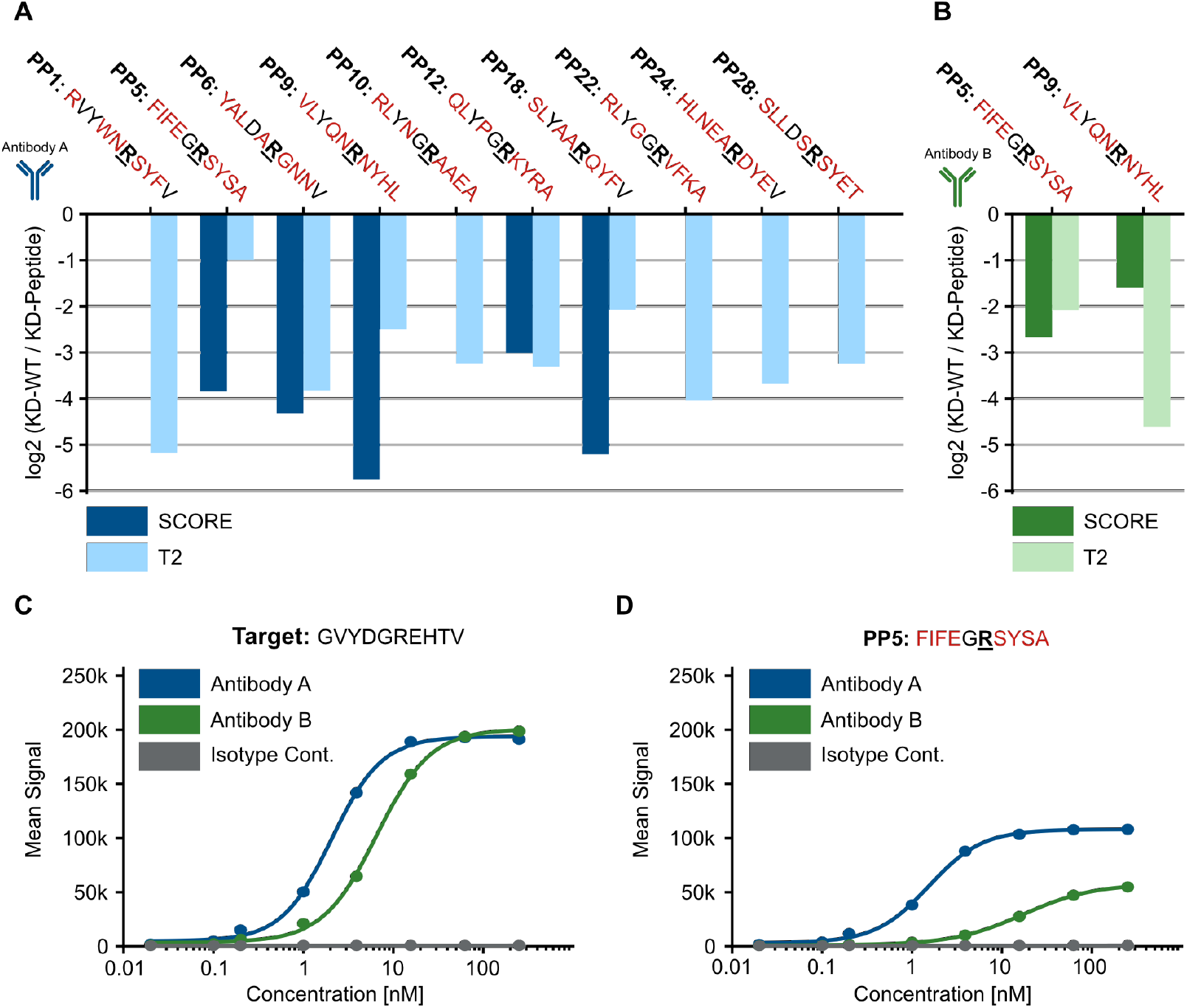
Validation of Predicted Off-Target Peptide Binding in SCORE microarrays and the T2 cell assay. (A) log2 binding ratios of SCORE and T2 binding experiments for Antibody A. (B) log2 binding ratios of SCORE and T2 binding experiments for Antibody B. No bars is equal to no binding detected. The more negative the values are, the weaker the binding. Positive values would have meant a stronger binding against the off-target peptide compared to the target. (C) *EC*_50_ dose–response titration of antibody A and antibody B on T2 cells loaded with the target peptide. (D) *EC*_50_ dose–response titration of antibody A and antibody B on T2 cells loaded with peptide PP5.

**Figure 5.**
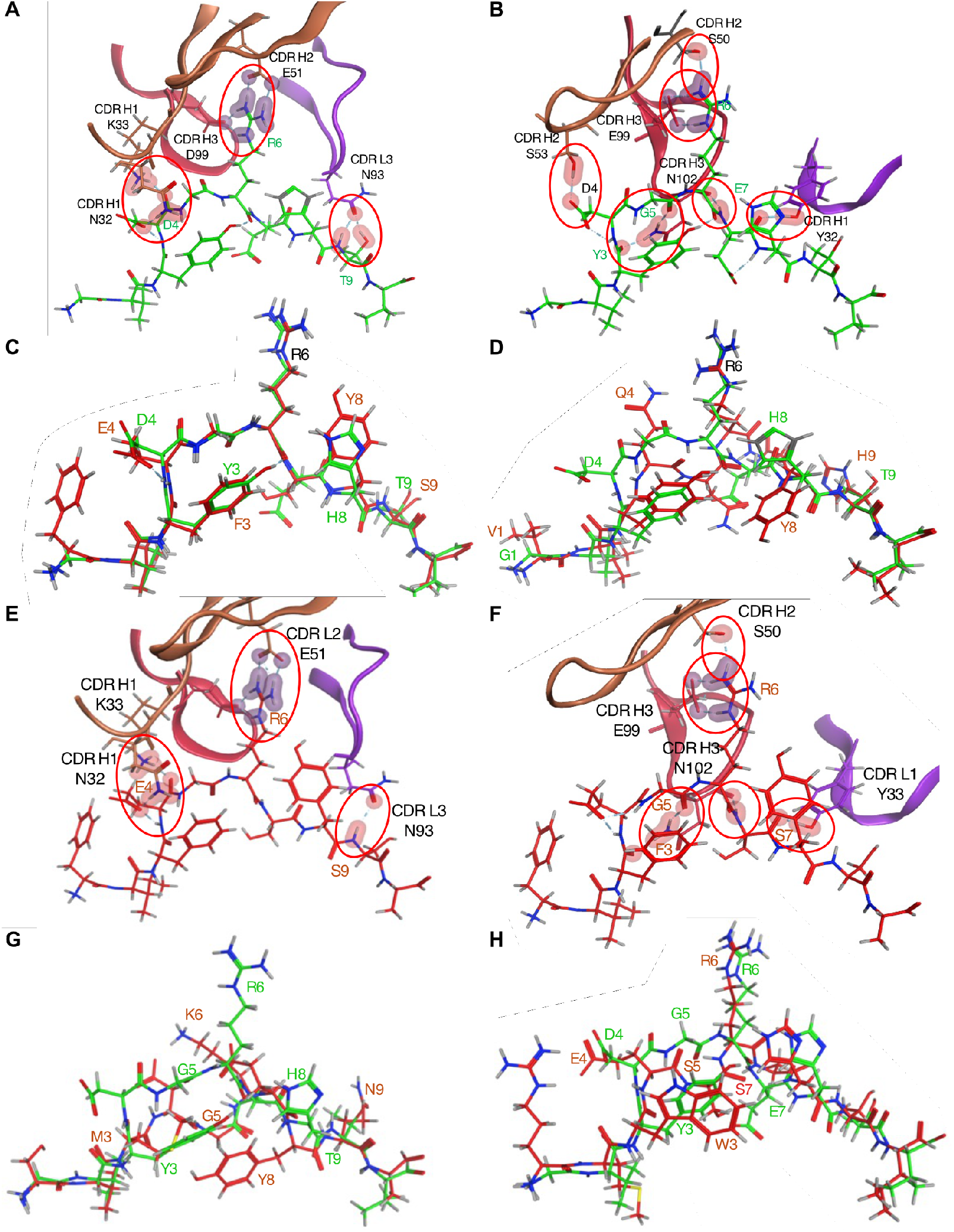
Structural basis of antibody-specific binding to predicted off-target peptides. (A) TCRm-MAGE-A4 peptide interfaces for antibody A as modelled by Chai-1 (F2). (B) TCRm-MAGE-A4 peptide interfaces for antibody B as modelled by Chai-1 (F2). Portions of the antibody and HLA-A*02:01 complex are omitted for clarity. Hydrogen bonds are shown as red clouds, ionic contacts as purple clouds. (C) Structural alignment of the MAGE-A4 (green) with PP5 (red) within the HLA-A*02:01 binding groove. (D) Structural alignment of the MAGE-A4 (green) with PP9 peptides (red) within the HLA-A*02:01 binding groove. Antibody and HLA complex are omitted for clarity. (E) Antibody A - PP5 interfaces. (F) Antibody B - PP5 interfaces. Hydrogen bonds are depicted as red clouds and ionic contacts as purple clouds. (G) Structural alignment of MAGE-A4 (green) with PP25 (red) in the HLA-A*02:01 groove. (H) Structural alignment of MAGE-A4 (green) with PP14 (red) in the HLA-A*02:01 groove. Antibody and HLA complex are omitted for clarity.

Both EpiPredict models identified off-target peptides with up to 8 mismatches relative to the WT sequence that still showed specific binding to TCRms. This demonstrates the model’s ability to capture complex, TCRm-specific binding determinants and to discover interactions with sequence-divergent, previously uncharacterized peptides highlighting its utility for off-target risk assessment and peptide discovery.

To assess peptide recognition in a cellular context, T2-cell binding assays were performed using the combined predicted off-target peptide set (Figure 4, A, B). T2 cells were pulsed with 100 µM of each peptide, a concentration sufficient for efficient loading of HLA-A*02:01 molecules in these TAP-deficient cells, as confirmed by increased *β* 2-microglobulin surface levels for 28 of 29 peptides predicted (Supplementary Figure S1). Peptide-pulsed cells were incubated with antibody A or antibody B, followed by staining with fluorophore-conjugated secondary antibodies. Binding was quantified by flow cytometry. Peptides that showed binding in SCORE also showed positive a signal in the T2 assay. The strongest binding peptide in the T2 cell assays (PP5) was further characterized in more detail by performing a complete *EC*_50_ dose-response curves. Antibody A showed an *EC*_50_ value of 2.1 nM for the target and 1.5 nM for PP5. Antibody B showed an *EC*_50_ value of 6.5 nM for the target and 17.1 nM for PP5. These results confirm that both antibodies bind PP5 in a similar range than the actual target sequence. The Isotype control antibodies showed no detectable binding across all conditions, confirming the specificity of the observed TCRm-peptide interactions.

In summary, T2-cell binding assays confirmed that machine learning-predicted peptides are recognized by TCRms in a physiologically relevant setting. These results support the conclusion that binding is not merely an artifact of recombinant pHLA complexes used in the SCORE experiments. Notably, EpiPredict identified antibody-specific off-target peptides with minimal or no sequence homology to MAGE-A4 that are presented by HLA-A*02:01 in a cellular context, reinforcing its value for off-target prediction beyond conventional peptide-centric approaches.

### Structural characterization of peptide-TCRm interactions predicted by EpiPredict

To investigate off-target peptides predicted by EpiPredict, we examined structural binding preferences of TCRm antibodies A and B. Because EpiPredict maps peptide sequences to antibody-specific binding probabilities, its predictions likely encode latent features of the paratope–epitope interaction landscape.

Baseline structural models of both antibodies with the WT MAGE-A4 peptide established their shared TCR-like binding mode: CDR-L3 and CDR-H3 dominated peptide contacts, while other CDR loops anchored the HLA helices (Supplementary Figure S2). Both antibodies required Arg6 for binding yet adopted distinct paratope conformations. Antibody A recognition involved CDR-L3 N93 to T9 and CDR-H1 N32/K33 to D4, reinforced by ionic interactions of CDR-H2 E51 and CDR-H3 D99 with R6. By contrast, antibody B relied on a dominant CDR-H3 E99-R6 salt bridge, with additional hydrogen bonds from CDR-H1 Y33 to E7, CDR-H3 N102 to Y3/G5/E7, CDR-H2 S53 to D4, and from CDR-H2 S50 to R6.

Consistent with their distinct engagement modes, antibody A recognized five predicted peptides, whereas antibody B bound only PP5 and PP9, which were also validated for antibody A. Despite substantial sequence divergence from MAGE-A4 (Hamming distances: 8 for PP5 and 8 for PP9) and strong differences between each other (Hamming distance PP5 vs. PP9: 8), both peptides adopted similar backbone conformations, exposing the conserved R6 (Figure 6b). This structural mimicry explains the observed shared cross-reactivity. For example, in PP5, antibody A retained strong ionic interactions with R6, complemented by hydrogen bonds to E4 and S9, whereas antibody B preserved its hallmark CDR-H3 E99-R6 salt bridge along with hydrogen bonds to F3, G5, and S7 (Figure 6c). In contrast, peptides lacking an exposed arginine for example PP25 (R6→K) failed to bind despite ML prediction. Structural modelling showed backbone displacement (>7 Å at G5), burying Lys6 too deep in the groove for antibody A CDR-H2 E51 and antibody B CDR-H3 D99 to engage, leaving only a weak H1–D4 contact (not shown). Similarly, PP14 maintained R6 but diverged structurally at P3-P7 (Figure 6d); absence of Y3 repositions R6 and attenuates its ionic interactions with CDR-H2 E51 and CDR-H3 D99. In addition, steric clashes between CDR-H1 and peptide W3/E4 precluded binding (not shown).

Together, these analyses identify Arg6 as a central determinant of TCRm binding and highlight how structure-based modelling contextualizes ML predictions. Importantly, EpiPredict uncovered off-target peptides with little sequence similarity yet structurally compatible features, demonstrating its utility for predicting paratope-epitope interactions beyond sequence homology. These insights may further guide optimization of TCRm specificity and inform safety assessments in therapeutic development.

## DISCUSSION

Comprehensive identification of potential off-target peptide–HLA complexes is a critical step in the development of TCR-mimic (TCRm) antibodies, as unintended cross-reactivity can cause severe toxicities [13, 14]. Conventional preclinical approaches such as alanine or X-scan mutagenesis [15, 16] provide early insights into residues critical for binding but are limited to sequence-centric consensus motifs, which often fail to capture sequence-dissimilar off-targets [9]. Structural analyses of TCRm–pHLA complexes [17] and high-throughput screening methods such as PresentER [18] or immunopeptidomics-based mass spectrometry [19] expand the range of detectable off-targets, yet each remains constrained by technical or biological context, including tissue specificity and peptide isolation biases [20, 21]. Together, these limitations highlight the need for scalable strategies that can interrogate the vast peptide–HLA sequence space and capture structural as well as sequence-based determinants of recognition.

Machine learning frameworks are well suited for this challenge, provided they have access to high-quality, domain-specific data. Building on this rationale, we developed EpiPredict, a TCRm-centric model trained on kinetic data from X-scan mutagenesis and sequence similar peptides. It is important to highlight, that the quality of this training dataset was key in order to train the antibody specific models for antibody A and B. Usually, high-throughput characterizations for TCRms are performed using MS-based methods. While these are great to capture potential off-targets, they are lacking true negative datapoints that are essential to train a model that can be used for classification tasks.

Trained on this data, EpiPredict, a small neural-network, predicted and validated off-target peptides that differed substantially from the cognate MAGE-A4 epitope. Despite the original training data only containing binders with an edit distance of up to 5, the models predicted only peptides with distances 6 or higher to be binders (Supplementary Table 1). Out of fold precision scores on the training data resulted in scores of 0.42 for antibody A and 0.30 for B. Upon validation, the precision of the models on the predicted peptide set dropped to 0.21 (3/14 predictions) for antibody A and 0.11 (2/18 predictions) for antibody B. These lower hit rates represent a reasonable performance given the inherent difficulty of predicting binding affinity for peptides with such high sequence dissimilarity from the training data. The ability to identify any true positives among highly divergent sequences (edit distance ≥6) demonstrates the model’s capacity to capture subtle binding patterns beyond simple sequence homology.

We propose that incorporating additional sequence positional context and 3D structural features may significantly improve predictive performance. Integrating physicochemical molecular surface descriptors of peptides, such as electrostatics and shape complementarity, may enhance EpiPredict’s ability to discriminate true binders. Recent deep learning frameworks have shown that extracting multi-dimensional structural features from predicted peptide–HLA conformations improves neoantigen immunogenicity prediction relative to sequence-only methods [22, 23], supporting their potential value for TCRm off-target modelling.

Similarly, embeddings from large protein language models [24, 25, 26] capture biologically meaningful sequence patterns and can serve as foundation representations for interaction models [27]. For example, TCR-ESM, a deep learning model, uses embeddings from ESM-1v (Evolutionary Scale Model-1v; H4) to predict TCR-peptide-MHC interactions with high accuracy [26]. The model encodes peptide, MHC, and TCR (CDR3*α*/CDR3*β*) sequences using ESM-1v and feeds them into a feedforward neural network to estimate TCR-pMHC interaction probabilities. Training is performed using both positive (experimentally validated binding pairs) and negative (mismatched) datasets, an approach similar to that used in EpiPredict. It is thus conceivable that EpiPredict could also be further enhanced by incorporating embeddings from large protein language models as a foundational representation of TCRm-pHLA interactions. These embeddings capture biologically meaningful patterns that may be difficult to encode using traditional sequence-based features, such as BLOSUM62-based encodings. Fine-tuning them on TCRm-pHLA-specific data could improve the model’s ability to learn interaction-relevant sequence features, resulting in more accurate off-target binding predictions.

Looking ahead, we plan to enhance EpiPredict by integrating it with EpiTox, an upcoming ML-based platform for comprehensive off-target risk assessment (manuscript in preparation). EpiTox will combine multiple data layers, including pHLA affinity, tissue-specific expression, and mass spectrometry validation, into a unified predictive framework. This integration will allow more precise evaluation of EpiPredict-identified peptides, supporting risk stratification and guiding TCRm optimization during preclinical development. Moreover, combining SCORE-based high-throughput pHLA screening with ML-driven prediction and structural validation may enable the de novo design of highly specific antibody-based TCRms for therapeutic applications, as has recently been demonstrated for simpler *α*-helical bundle scaffolds [28] an approach that is now actively pursued in our digital laboratories.

In summary, our study demonstrates that EpiPredict can effectively identify structurally mimetic, sequence-dissimilar off-target peptides in a TCRm-centric context. Importantly, many predictions were experimentally validated as true binders using both SCORE-based interaction assays and T2-cell binding assays. While limitations remain, especially in capturing strict positional requirements, incorporating structural and language-model-derived representations offers promising routes for improvement. Our findings position EpiPredict as a valuable tool for early-stage assessment of off-target risk across the peptidome, extending beyond traditional sequence-similarity approaches.

## MATERIALS AND METHODS

### Identification of fully human TCR mimic anti–MAGE-A4 antibodies

Fully human TCR-mimic antibodies targeting MAGE-A4 were generated in collaboration with Biocytogen Pharmaceuticals Co., Ltd. (Beijing, China), using their RenMice-based TCR-mimic antibody platform, as previously described [29]. In this system, genetically engineered RenMice co-express a fully human antibody repertoire alongside HLA-A*02:01 and were immunized with recombinant MAGE-A4 peptide-HLA-A*02:01 complexes. B cells from immunized mice were subsequently screened using the Beacon on-chip platform to enable high-throughput discovery of TCR-mimic antibodies. Two highly specific anti-MAGE-A4 TCR-mimic antibodies, antibody A and antibody B, were identified, demonstrating superior functional and biophysical properties.

### Antibody and Fab Production

Recombinant full-length antibodies (Antibody A, antibody B, and the isotype control Palivizumab, incorporating a human IgG1 Fc LALA-PG mutation [30] and Fab fragments of antibody A and B (C-terminal 3×Myc-1×His8 tag) were produced in collaboration with Biointron (Beijing, China). Heavy and light chain sequences were synthesized and cloned into pCDNA3.4 expression vectors, followed by co-transfection into CHO-K1 cells. Proteins were expressed in suspension culture for 4–6 days at 37°C, 120 rpm, and 8% CO_2_. IgG antibodies were purified via Protein A affinity chromatography (MabSelect PrismA/MabSelect Sure; Cytiva Life Sciences, Marlborough, Massachusetts, USA), eluted with sodium acetate (pH 3.4), and dialyzed into PBS (pH 7.2–7.4). Fab fragments were purified using Ni-NTA resin under standard conditions. Protein purity, monomeric state, and integrity were assessed by analytical size-exclusion chromatography (SEC).

### Peptide–HLA array production and SCORE measurements

Peptide–HLA array production and SCORE measurements were performed essentially as described previously [31]. Peptides were purchased from Peptides & Elephants GmbH (Hennigsdorf, Germany), and biotinylated, stabilized HLA-A*02:01 variant was used [32]. Antibody A and B were measured in their fab format to avoid avidity effects. The binding curves were fitted and analyzed using an internal version of anabel and a 1:1 Langmuir binding model [33].

### BLI Binding Assay

BLI experiments were performed on an Octet R8 system (Sartorius, Göttingen, Germany). Biotinylated human HLA-A*02:01 & *β*_2_M & MAGE-A4 (GVYDGREHTV) complex protein (Acro Biosystems, HLM-H82E5) was immobilized at 30 nM on Streptavidin (SA) Biosensors. Analytes and ligands were diluted in kinetics buffer (1× DPBS, 0.1% casein, 0.05% Tween-20). Biosensors were hydrated in kinetics buffer for 10 min prior to use. Assays were conducted at 25^*°*^C with shaking at 1000 rpm. Sensors were equilibrated in kinetics buffer for 60 s, loaded with pHLA for 200 s, followed by a 60s wash step, 120 s baseline, a 350 s association, and a 600 s dissociation phase. Three Fab concentrations (60, 20, and 6.7 nM) were tested. A pHLA-loaded sensor incubated in buffer served as a reference for subtraction. Data were analyzed using Octet Analysis Studio 13.0. Sensorgrams were aligned to the end of the baseline step, and inter-step correction was applied to match the start of the dissociation phase. Data were smoothed using Savitzky–Golay filtering and globally fitted with a 1:1 Langmuir binding model.

### T2 cell binding assay

T2 cells (DSMZ# ACC598) were cultivated at 37^*°*^C and 5% CO2 in RPMI-1640 medium (Thermo Fisher Scientific, Waltham. USA) supplemented with 10% heat-inactivated FCS. For peptide pulsing, peptide solution and T2 cells were mixed in 96-multiwell plates directly or pre-mixed in falcon tubes using AIM-V medium (Thermo Fisher Scientific). Peptide pulsing happened over night for 17-18 hours at 37°C and 5% CO2. The next day the multi-well plate was centrifuged at 700 × g for 1 min at 4°C. The supernatant was discarded. Cells were washed once with PBS and LIVE/DEAD Fixable Violet stain (Thermo Fisher Scientific, 1:1000 dilution in PBS) was added. The cells were re-suspended and incubated on ice for 30 min with gentle shaking, protected from light. After staining, 170 *µ*l of PBS was added per well, followed by centrifugation (1 min at 700 × g, 4°C). The supernatant was discarded, and cells were washed twice with 200 *µ*l of FACS buffer (PBS + 2% FCS). For peptide pulsing control, an anti-B2M-FITC antibody (Sigma-Aldrich, Burlington, USA, #SAB4700012) or anti-B2M-AF647 antibody (Thermo Fisher Scientific, #MA5-18119) was diluted 1:225 in FACS buffer and added to the wells. The cells were resuspended and incubated for 30 min on ice with gentle shaking. TCRm binding analysis was either performed with antibody A and B in IgG format in single dose measurement or full dose 8-point titration. After pulsing, cells were spun down, supernatant was removed and TCRm antibody dilutions were added. This was followed by gently resuspension and incubation on ice for 1h and gentle shaking. After incubation, cells were washed once with FACS-buffer and secondary anti-Fc-detection antibody was added (Thermo Fisher, #A18818). This was followed by gently resuspension and incubation on ice for 1h and gentle shaking. Subsequently, cells were washed twice with FACS-buffer. Flow cytometry was performed using the Attune NxT flow cytometer (Thermo Fisher). Data analysis was performed with FlowJo (BD Life Sciences, Franklin Lakes, USA). Gating strategy for all experiments comprised pulse geometry gating and Live/Dead Gating. Compensation, if required, was done with FlowJo. Data visualization was done with GraphPad Prism (Dotmatics, Boston, USA).

### Labeled kinetic dataset generation

To generate antibody-specific training data for EpiPredict, we employed a stepwise workflow beginning with in silico off-target prediction using the EpiTox pipeline (manuscript in preparation). EpiTox identified candidate peptide sequences from the human proteome with up to five substitutions relative to the cognate MAGE-A4 epitope (GVYDGREHTV), filtered for predicted HLA-A*02:01 presentation and expression relevance. These candidate sequences were combined with the complete single-amino acid substitutional scan (X-scan) of the wild-type peptide, yielding a set of 530 unique peptides.

Custom peptide-HLA SCORE sensorchips (microarrays) were generated from this panel as described above, and binding interactions with antibody A and B were quantified using SCORE. Raw sensorgrams were processed with the anabel [33] analysis framework, which performed automated curve fitting and parameter extraction for kinetic association (*k*_*on*_), dissociation (*k*_*o f f*_), and affinity (*K*_*D*_). Rule-based quality control was subsequently applied using in-house software. Complexes with *K*_*D*_ values reproducible across independent replicates were designated as binders; all others were classified as non-binders. The resulting datasets (530 peptides per antibody) were curated and stored, establishing a labeled kinetic dataset that served as direct input for training the EpiPredict neural network models.

### Unlabeled 10mer peptide dataset generation

To extend predictions beyond the experimentally profiled panel, we generated a comprehensive in silico peptide space representing potential HLA-A*02:01 ligands across the human proteome. All canonical and isoform sequences from UniProt (release February 2025) totaling 34,184 protein sequences were parsed with a sliding window of 10 amino acids (stride = 1), yielding a library of >10 million unique 10-mers.

This library was filtered for predicted HLA-A*02:01 presentation using NetMHCpan 4.1 [11], applying a rank threshold of ≤2 to select peptides with high likelihood of stable MHC binding. Redundant sequences and low-confidence predictions were removed, resulting in a final set of 316,855 unique peptides. Unlike the labeled kinetic dataset, this in-silico collection was used without binding class annotations and served as the unlabeled prediction space for proteome-wide inference.

### EpiPredict model architecture and training

The neural network models were MultiLayer Perceptrons (MLPs) implemented in Python using the PyTorch framework. The MLP consisted of two layers with a ReLU activation in-between. A sigmoid activation was applied to the model output to produce probability scores for binary classification (binder /non-binder). Peptide sequences were encoded into fixed-size tensor representations using BLOSUM62 amino acid substitution matrices. Hyperparameters were optimized using five-fold cross validation with the binary cross-entropy loss function. To minimize performance variance across training iterations, at each fold an ensemble of five MLPs were trained, with each model’s weights initialized to different values. The models were trained using the Adam optimizer and the One-Cycle learning rate scheduler. The full set of training parameters and final selected hyperparameters are provided in Table 2. The training procedure resulted in a nested ensemble of 25 MLPs, five models at each fold for five folds. During inference, the binding probability was calculated as arithmetic mean across all models.

**Table 2.** Neural Network parameters and hyperparameters.

| Parameter | Value |
| --- | --- |
| Number of layers | 2 |
| Layer sizes | 200-16-1 |
| Number of models in ensemble | 5 per fold |
| Activation function | ReLU |
| Optimizer | Adam |
| Scheduler | Onecycle |
| Batch size | 8 |
| Max Learning rate | 0.01 |
| Number of epochs | 15 |
| Loss function | binary cross-entropy |

### Structure modelling with Chai-1 and MOE post-processing

Antibody-pHLA complexes for antibody A and antibody B with HLA-A02:01 presenting MAGE-A4(230-239) peptide were modeled with Chai-1 in single-sequence mode across a small grid of sampling parameters (recycles 3-20; diffusion steps 200-1000), with and without restraints. Based on recurrent features in TCRm-pHLA structures, we used targeted restraints: for antibody A, CDR-L2 E55 → HLA R65 and CDR-H2 E51 → peptide R6; for antibody B, CDR-H3 D109 →HLA R65 and CDR-L1 Y32 → peptide E7. All models were prepared in MOE (Quick-Prep/Protonate3D; standard Amber:EHT reaction-field settings), then minimized with backbone tethers to remove local clashes while preserving the pose.

Models were ranked using no-reference interface metrics after MOE relaxation: buried surface area (BSA), a shape-complementarity proxy, clash density, hydrogen bonds and salt bridges and ensemble stability (contact-map Jaccard after receptor superposition). We further computed interaction energies for antibody-pHLA (HLA+peptide) and antibody-peptide contacts and derived a peptide-preference index. Under these criteria, optimal sampling was 20 recycles/600 steps for the antibody A - pHLA complex and 20 recycles/800 steps for the antibody B - pHLA complex, both with restraints. Full protocols, parameter values, and normalization/ranking are provided in supplementary materials and methods.

### Databases

Our antibodies and molecules are fully tracked using the Genedata Biologics (GDB) system [34].

## Supporting information

experiment_raw_data

## AUTHOR CONTRIBUTIONS

A.S. and S.K. designed, developed, and applied the EpiPredict model. A.S., S.K., C.R., S.S., H.aH., O.S., and J.B. designed and implemented the experimental workflow. H.aH. performed the initial EpiTox-based off-target predictions. P.H. produced the SCORE microarrays. T.H. conducted SCORE and BLI-based measurements. J.S. carried out T2 cell-based experiments. S.S. designed and supervised analysis of cell-based experiments. C.R. performed structural modelling and analyses. J.B. wrote the manuscript. All authors contributed to manuscript review and revisions.

## DATA AND CODE AVAILABILITY

The EpiPredict code is available from the corresponding author upon reasonable request for academic or non-commercial research purposes. The raw data used used in this study is available in the supplement.

## ACKNOWLEDGMENTS

The authors gratefully acknowledge Martin Klatt (Charité Berlin) and Björn Steinmann (BioCopy AG) for their valuable contributions to this work. AI use: GPT-4, GPT-5 and Claude Sonnet 4 models were employed exclusively for language refinement during manuscript preparation.

## CONFLICTS OF INTEREST

All authors are current employees of BioCopy AG or BioCopy GmbH. They may hold shares or stock options in BioCopy AG. BioCopy has patent applications relating to certain research areas described in this article.

