## Supplementary_Figures for "Beyond Sequence Similarity: ML-Powered Identification of pHLA Off-Targets for TCR-Mimic Antibodies Using High Throughput Binding Kinetics"

**Figure S1**

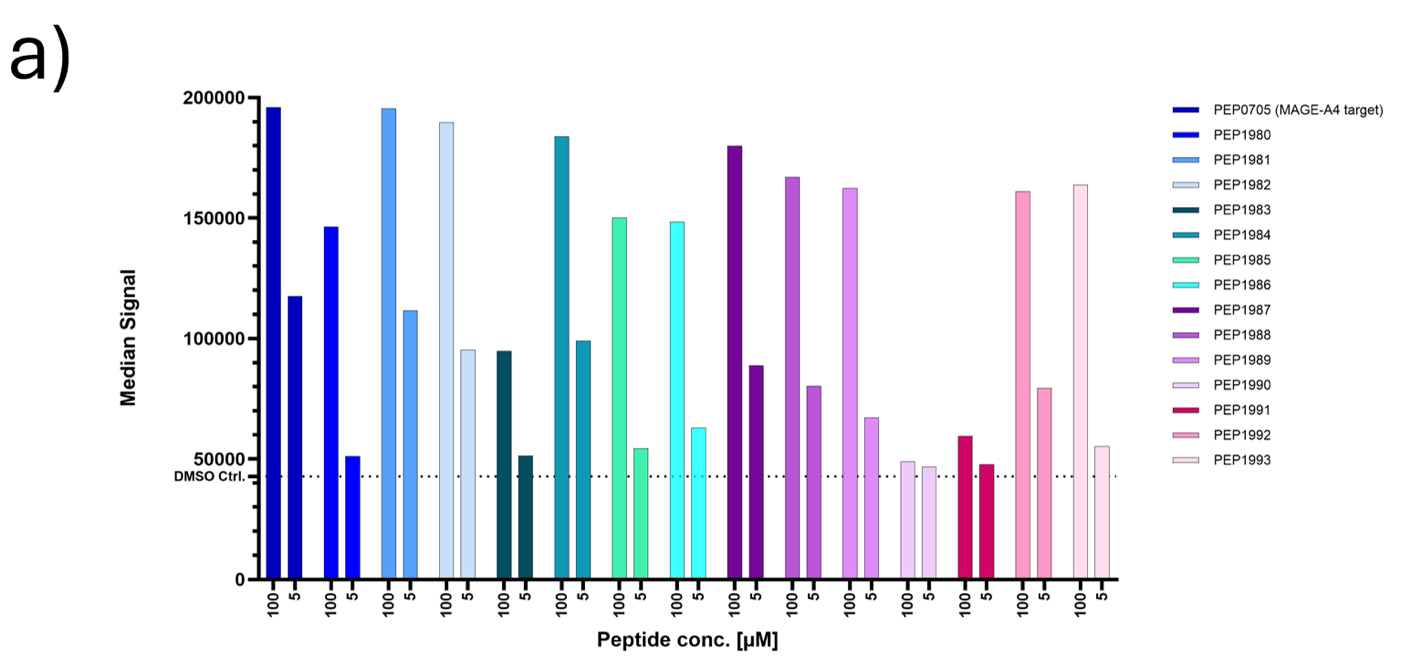

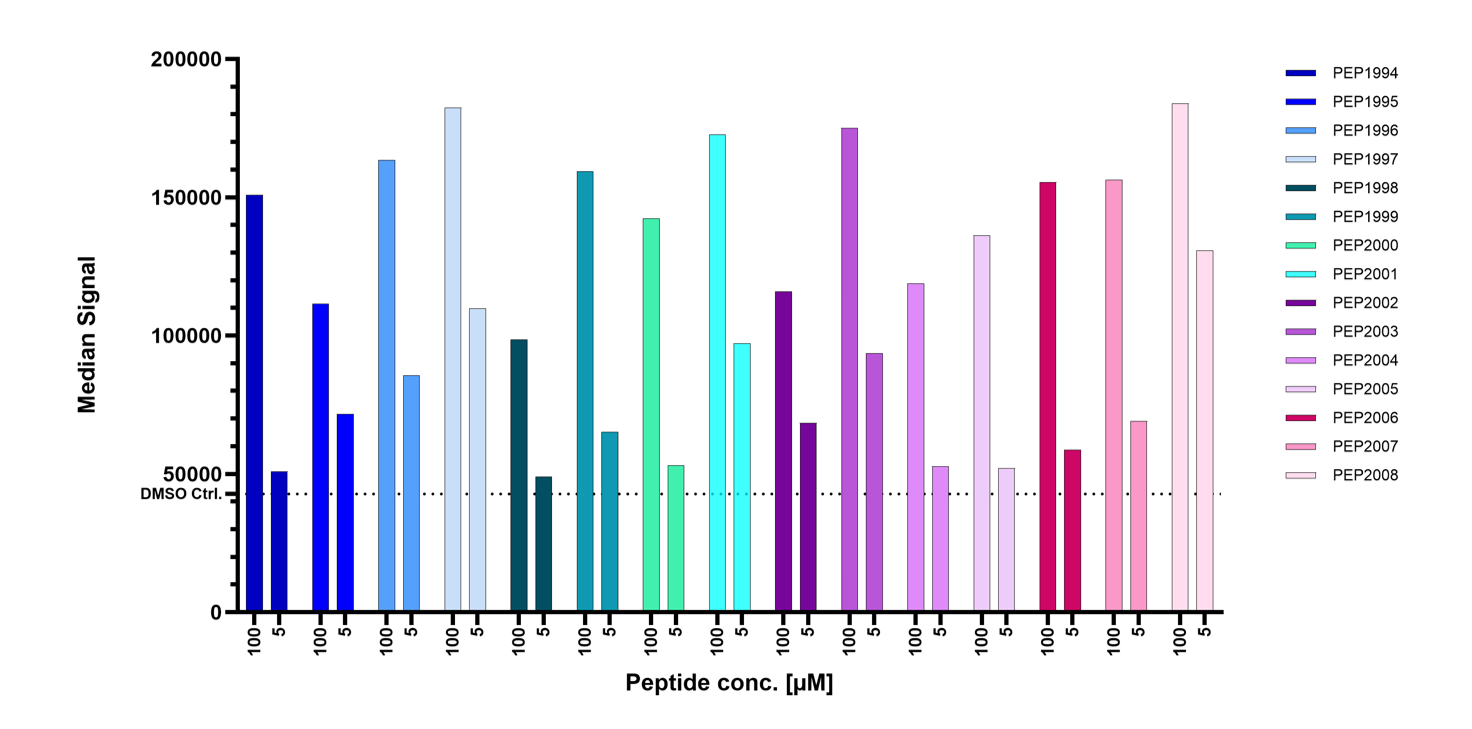
**A**

**B**

**Figure S2.** Validation of Predicted Off-Target Peptide Binding in the T2 cell assay. **(A)** T2 cells were loaded with 100 µM or 5 µM of either the wild-type decapeptide or off-target peptides predicted by EpiPredict. To control for false negatives due to insufficient peptide presentation, peptide loading was verified by β₂-microglobulin staining. Peptide PEP1990 could not be loaded even at 100 µM.

**Figure S2**

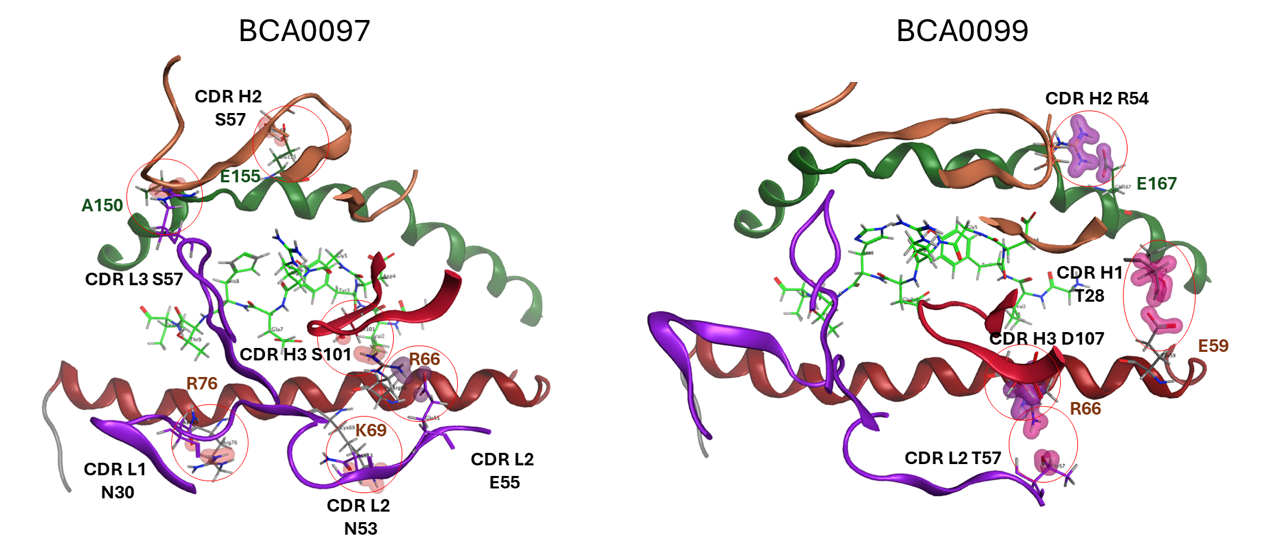

**Figure S2.** Structural basis of antibody-specific binding to predicted off-target peptides.

Hydrogen bonds are shown as red clouds and ionic contacts as purple clouds. The HLA α1-helix is colored red, and the α2-helix green. Chai1 modelling indicates that BCA0097 binds the pHLA complex in a TCR-like orientation, anchored by six principal interactions: hydrogen bonds between CDR-L1 N30 and R76 (α1-helix), CDR-L2 N53 and K69 (α1-helix), CDR-L3 R96 and A150 (α2-helix), CDR-H2 S57 and E155 (α2-helix), CDR-H3 S101 and R66 (α1-helix), and an ionic contacts between CDR-L2 E55 and R66 (α1-helix). The BCA0099 model reveals strong ionic interactions anchoring the antibody centrally in a TCR-like orientation: CDR-H2 R54 to E167 (α2-helix) and CDR-H3 D107 to R66 (α1-helix), complemented by a network of hydrogen bonds involving CDR-H1 T28 to E59 (α1-helix), CDR-H3 W101 to K67 (α1-helix), and CDR-L2 T57 to R66 (α1-helix).
