## Supplementary_Material_and_Methods for "Beyond Sequence Similarity: ML-Powered Identification of pHLA Off-Targets for TCR-Mimic Antibodies Using High Throughput Binding Kinetics"

**Structure modelling with Chai-1 and MOE post-processing**

Antibody-pHLA complexes for antibody A and antibody B with HLA-A*02:01 MAGE-A4 (230 239) were modeled using Chai-1, a multimodal foundation model for biomolecular structure prediction that supports single-sequence inputs and optional constraint features (pocket/contact/docking) to encode prior knowledge about interfaces (14). We ran Chai‑1 in single‑sequence mode and generated ensembles per complex across a grid of sampling parameters (recycles: 3-20; diffusion steps: 200, 400, 600, 800, 1000) with and without restraints. Generated models were carried forward to MOE (Molecular Operating Environment, Chemical Computing Group, version 2024.0601) for structure preparation prior evaluation with no‑reference interface metrics (see below).

To define a structural baseline and peptide pose, modeled pHLA coordinates (MHC heavy chain + β2m + peptide) were superposed to the cryo‑EM structure 8FJA (HLA‑A*02:01 presenting MAGE‑A4 230-239 bound by REGN6972) using an MHC+peptide alignment.

Because several TCRm-pHLA structures (PDB: 7BBG, 8FJA, 3GJF, 7RE7) frequently show an acidic residue (Asp/Glu) in CDR‑L2 (positions 50-55) forming a strong ionic interaction with HLA helix arginine R65, we tested antibody A CDR‑L2 D50 and E55 as anchor restraints. Among these candidates, the E55(L2) → R65(HLA) contact produced the most consistent and favorable poses. By analyzing top ranked antibody A models, a second anchor point was proposed (H2 E51 → peptide R6). The combination of CDR‑L2 E55→R65 (HLA helix) and CDR‑H2 E51→R6 (peptide) restraints was subsequently used for modeling antibody A -pHLA complexes. Based on structure from 8FJA and high ranked models from antibody A we proposed for antibody B contact points H3 D109 -> R65 (HLA helix) and L1 Y32 to E7 (peptide). Both were used for modeling antibody AB-pHLA complexes.

Sampling settings were selected based on ensemble stability and interface quality criteria after MOE relaxation: high shape‑complementarity proxy (Sc_proxy), adequate buried surface area (BSA), low clash density (per 1000 Å²), strong peptide‑centric interaction energy, and contact‑map stability (Jaccard) across seeds. Under these criteria, antibody A performed best at 20 recycles and 600 diffusion steps (with restraints), whereas antibody B performed best at 20 recycles and 800 diffusion steps (with restraints).

For each modeled complex, MOE was used for structure preparation. QuickPrep -> preserve sequence and neutralize; Protonate3D with ASN/GLN/HIS flips; remove waters farther than 4.5 Å from receptor or ligand; enforce standard residues. Backbone restraints: receptor (pHLA) backbone atoms (N, CA, C, O) strongly tethered (strength 10, buffer 0.25); antibody backbone moderately tethered; side chains free. Energy minimization: AmberEHT with bonded, van der Waals, electrostatics, and restraints enabled; non‑bonded cutoff on/off 8/10 Å; reaction‑field dielectric with interior ε=1 and exterior ε=80. Two short stages: (i) hydrogens + side chains, (ii) all atoms with backbones tethered. With tethered backbone atoms, we used the minimization algorithm.

**Analysis and ranking of Chai-1-predicted antibody-pHLA structures**

PDB files were parsed with Biopython; chain selections defined receptor (pHLA) and ligand (antibody). For pairwise comparison, the moving model was superposed onto the fixed model by fitting shared receptor CA atoms (≥3 shared positions), and the transformation was applied to all atoms prior to interface calculations.

Residue-residue contacts were defined between heavy atoms across partners with a 5.0 Å cutoff. To remove dependence on chain IDs/numbering, canonical keys were built by chain‑length rank and residue order; these yielded interface residue sets and a contact‑pair set for each model. Pose similarity was quantified by the Jaccard index of these contact sets.

Buried surface area (BSA): SASA(Receptor) + SASA(Ligand) − SASA(Complex[R∪L]) using FreeSASA (Ref. 42; Mitternacht 2016) after removing waters/ions and selecting only receptor/ligand chains (Å²). Shape‑complementarity proxy (Sc_proxy): nearest‑neighbor cross‑partner distances with an interface cutoff at 4.5 Å, upper‑tail trimming (30%), mapped to 1/(1+(d_mean/3.0)^2) in [0,1]. Hydrogen bonds: cross‑partner N⋯O heavy‑atom distance ≤ 3.5 Å. Salt bridges: Lys NZ / Arg NH1,NH2 to Asp OD1,OD2 / Glu OE1,OE2 ≤ 4.0 Å. Cβ contacts: count of Cβ-Cβ (Gly uses CA) pairs ≤ 8.0 Å across partners. Clash density: clashes per 1000 Å² BSA where a clash is distance < r_vdW(A)+r_vdW(B)−0.6 Å (element‑specific radii).

To summarize pose families without a native reference, we computed (i) the Jaccard similarity (Ref. 43; Martin 2022) between canonical contact sets and (ii) a ligand‑interface iRMSD to a base model. The base model was chosen among top‑scoring models as the structure minimizing 0.6·(1−Jaccard) + 0.4·[1/(1+(iRMSD/2.0)^2)] to all others. Ligand iRMSD was computed on shared CA positions of interface residues after receptor superposition.

Interaction energies (kcal·mol⁻¹) were calculated (MOE) for Antibody-(HLA+peptide) and antibody-peptide contacts. Energies were treated by magnitude (more negative = stronger) and normalized to [0,1] using empirical 10th-90th percentiles of absolute values. A peptide‑preference index r = [E_Ab-peptide] / [E_Ab-(HLA+peptide)] (clamped to [0,1]) quantified peptide‑dominated binding. Absolute energy values from the fixed MOE setup are interpreted as relative metrics.

Normalization and ranking were performed as follows. BSA_norm: linear map with anchors 700 Å² -> 0 and 2000 Å² -> 1. Sc_norm: Sc_proxy clamped to [0,1]. HB/SB/CB_norm: saturation x/(x+K) on densities per 1000 Å² BSA with K = 3.0 (H‑bonds), 1.0 (salt bridges), 25.0 (Cβ). Clash_norm: decreasing 1/(1+x/2.0) on clash density (per 1000 Å² BSA). Energy normalization and peptide‑preference: as defined above.

We used a peptide‑focused scoring scheme that upweights Ab-peptide energy and the peptide‑preference index while retaining structural gates, with weights renormalized to available features: a no‑reference heuristic combining Sc_norm, BSA_norm, HB/SB/CB_norm and Clash_norm. Gate penalties (×0.5 each) were applied if BSA<700-800 Å², Sc_proxy<0.50, or clash density>3-4 per 1000 Å². Final tables were sorted by score descending.

All computations used custom Python (Biopython (Ref. 44; Cock 2009) for parsing/superposition; FreeSASA for SASA/BSA; NumPy (Ref. 45; Harris 2020)/pandas (Ref. 46; McKinney 2010) for data processing; Streamlit UI). The app exports raw metrics, normalized features, and ranked tables (CSV) to enable reproduction of rankings from primary features.

Results for top 20 models (antibody A and antibody B) are shown in Table 1 and 2 below.

**Table 1**

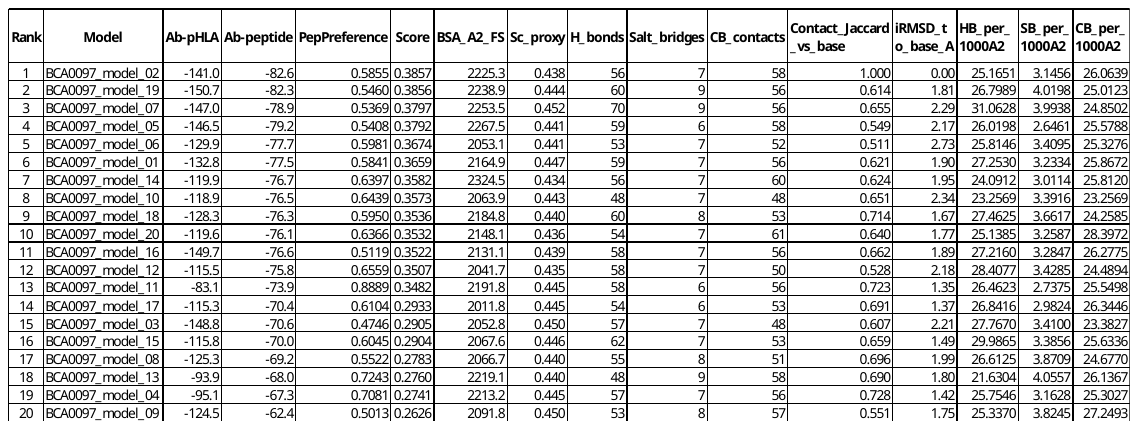

**Table 2**

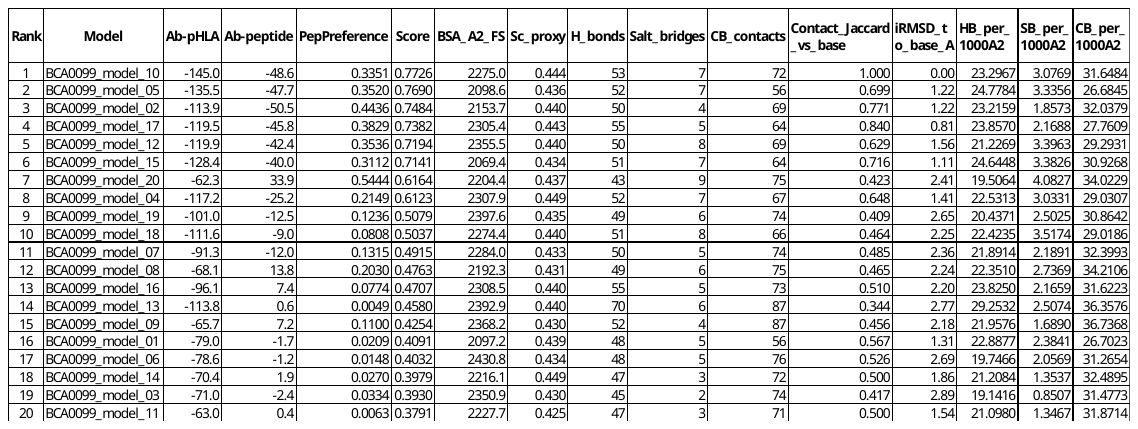

**Table 1 and 2: Ranking of top 20 Chai-1 antibody-pHLA models.**

Models of BCA0097 (Antibody A) / BCA0099 (Antibody B) bound to HLA-A*02:01-MAGE-A4 (230-239) were prepared with MOE (QuickPrep, Protonate3D, restrained minimization) and evaluated with our pipeline (no reference). The table reports, per model, structural interface metrics (buried surface area, shape-complementarity proxy, hydrogen bonds, salt bridges, Cβ-Cβ contacts), ensemble stability metrics relative to the selected base model (contact Jaccard index; ligand-interface iRMSD), and interaction energies (kcal·mol⁻¹) computed in the fixed AmberEHT setup for the antibody vs. pHLA complex (Ab-pHLA) and for the antibody vs. peptide (Ab-peptide). Ranking is peptide-focused: more favorable (more negative) Ab-peptide energies and stronger peptide preference contributed most to the final score, while structural metrics acted as consistency checks; clash metrics and all internal normalized features were omitted for compactness. Lower iRMSD and higher Jaccard indicate better agreement with the base model (for BCA0097: BCA0097_model_02; for BCA0099: BCA0099_model_10). All energies are relative within the stated force-field and cutoff settings.

Ab-pHLA, Ab-peptide: MOE (AmberEHT, RF ε = 1/80, 8/10 Å cutoff) interaction energies (kcal·mol⁻¹); more negative = stronger. PepPreference: (Ab-peptide) / (Ab-pHLA) clamped to [0,1] (larger = more peptide-dominated). Score: peptide-focused composite (weights renormalized to present features: Epep_norm: 0.35; PepPreference: 0.20; Sc_norm: 0.20; BSA_norm: 0.10; Clash_norm: 0.05; Ecomplex_norm: 0.03; HB_norm: 0.03; SB_norm: 0.02; CB_norm: 0.02). Sorting: by Score (descending). BSA_A2_FS: buried surface area (Å²) via FreeSASA for selected chains; Sc_proxy: packing/shape-complementarity proxy in [0.1]. H_bonds, Salt_bridges, CB_contacts: cross-partner counts; corresponding _norm columns use saturation maps (K = 3.0, 1.0, 25.0). Contact_Jaccard_vs_base: canonical contact-map Jaccard vs. base; iRMSD_to_base_A: ligand-interface iRMSD (Å) to base model after receptor fit. BSA/Sc/Clash_flag: gatekeeping flags (BSA < 700-800 Å²; Sc < 0.50; clash density > 3-4 per 1000 Å²). HB/SB/CB_per_1000A2: densities per 1000 Å² BSA.
